# Chromosome-level genome assembly for the Aldabra giant tortoise enables insights into the genetic health of a threatened population

**DOI:** 10.1101/2022.04.20.488802

**Authors:** F.G. Çilingir, L. A’Bear, D. Hansen, L.R. Davis, N. Bunbury, A. Ozgul, D. Croll, C. Grossen

## Abstract

The Aldabra giant tortoise (*Aldabrachelys gigantea*) is one of only two giant tortoise species left in the world. The species is endemic to Aldabra Atoll in Seychelles and is considered vulnerable due to its limited distribution and threats posed by climate change. Genomic resources for *A. gigantea* are lacking, hampering conservation efforts focused on both wild and ex-situ populations. A high-quality genome would also open avenues to investigate the genetic basis of the exceptionally long lifespan. Here, we produced the first chromosome-level *de novo* genome assembly of *A. gigantea* using PacBio High-Fidelity sequencing and high-throughput chromosome conformation capture (Hi-C). We produced a 2.37 Gbp assembly with a scaffold N50 of 148.6 Mbp and a resolution into 26 chromosomes. RNAseq-assisted gene model prediction identified 23,953 protein-coding genes and 1.1 Gbp of repetitive sequences. Synteny analyses among turtle genomes revealed high levels of chromosomal collinearity even among distantly related taxa. We also performed a low-coverage re-sequencing of 30 individuals from wild populations and two zoo individuals. Our genome-wide population structure analyses detected genetic population structure in the wild and identified the most likely origin of the zoo-housed individuals. The high-quality chromosome-level reference genome for *A. gigantea* is one of the most complete turtle genomes available. It is a powerful tool to assess the population structure in the wild population and reveal the geographic origins of ex-situ individuals relevant for genetic diversity management and rewilding efforts.

## Background

As human activities drive our planet into the sixth mass extinction[1], genomic technologies are becoming an increasingly important tool for conservation researchers. The establishment of reference-quality genomes for species of conservation concern makes key contributions to the study of common genetic health issues elucidating the full spectrum of genomic diversity, accurately quantifying inbreeding, mutation load, and introgression, detecting hybridization, and identifying adaptive variation in the face of rapidly changing environments[2]. The number of available reference genomes for non-model species has been increasing thanks to ongoing efforts in several global genome consortia, such as the Earth Biogenome Project[3], the Vertebrate Genomes Project[4,5], and the Global Invertebrate Genomics Alliance[6]. However, available reference genomes of non-model species are not homogeneously distributed across the tree of life. Only three reference genomes represent the Testudinidae family (or tortoises) from two genera, with two genomes being annotated and only one assembled to chromosome level. Tortoises have been integral components of global ecosystems for about 220 million years[7] contributing to seed dispersal, nutrient and mineral cycling, and carbon storage[8]. Over their long evolutionary history, giant tortoises, in particular, have evolved a life history characterized by delayed maturity, extended reproductive lives, and extreme longevity[9].

Currently, there are only two extant giant tortoise taxa, both of which face extinction threats[10]. Galápagos giant tortoises (*Chelonoidis niger* and subspecies thereof, formerly *Chelonoidis niger* species complex) are native to the Galápagos Islands in the eastern Pacific Ocean, and taxa of this group are listed as either vulnerable, endangered, or extinct according to IUCN Red List (v2.3). Aldabra giant tortoises (*Aldabrachelys gigantea*) (Fig. 1A) are native to Aldabra Atoll in the western Indian Ocean (Fig. 1B). Due to their extremely limited distribution in the wild and the threats posed by climate change, the species is vulnerable. Genomes of giant tortoises may harbor clues to their exceptional life-history traits. Assessing genome-wide variation within species, including deleterious mutation load, will critically improve conservation management programs[11]. The recently established reference genome for one of the Galápagos giant tortoises, *Chelonoidis niger abingdonii*, revealed insights into potentially aging, disease, and cancer-related gene functions by analyzing gene content evolution among tortoises[12]. For Aldabra giant tortoises, only short-read sequencing data is available from the same study[12].

**Fig. 1.**
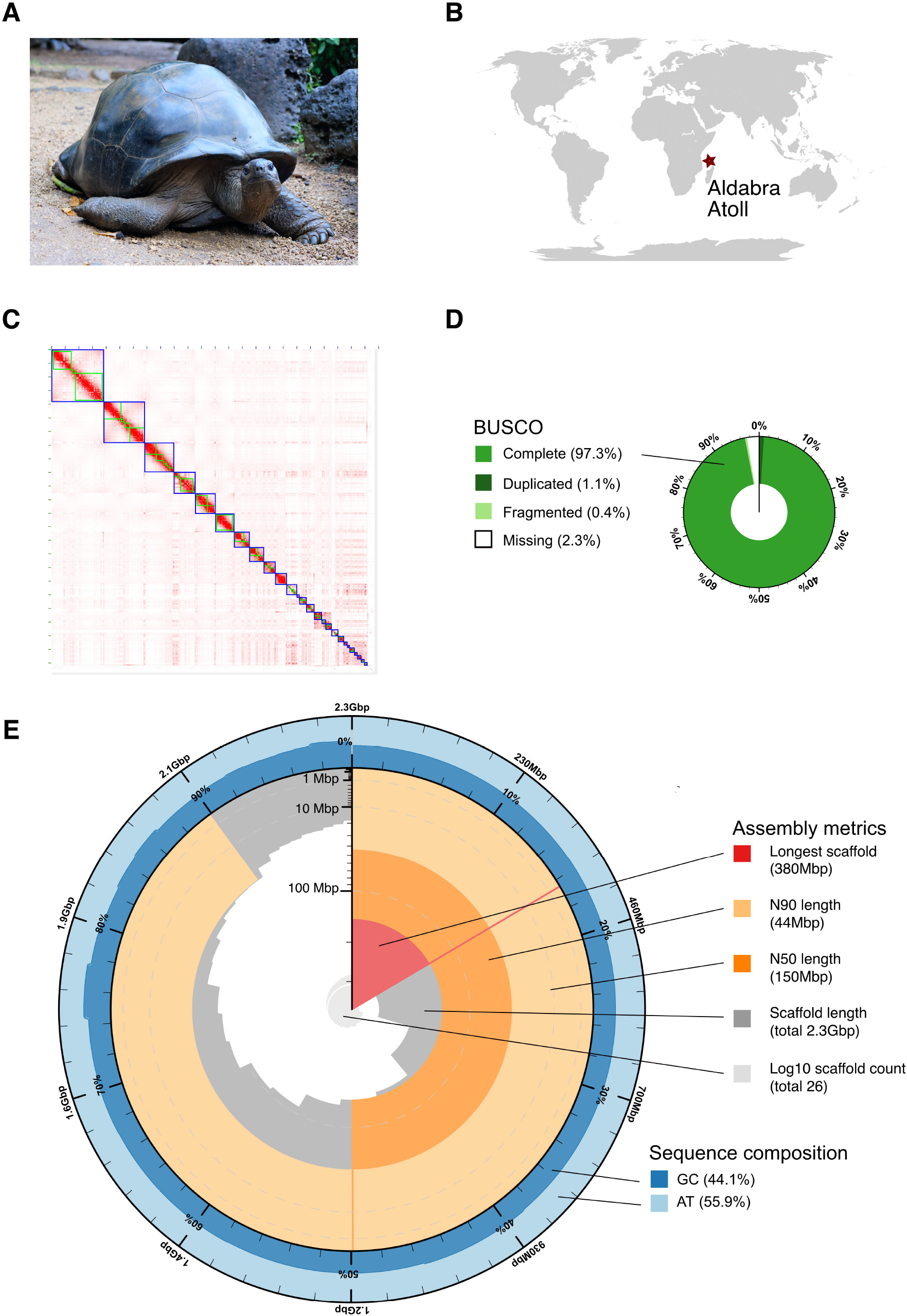
**A)** A female *Aldabrachelys gigantea* resting at La Vanille Nature Park, Mauritius. **B)** World map showing the location of Aldabra Atoll. **C)** Hi-C contact map of the chromosome-level assembled *Aldabrachelys gigantea* reference genome. Blue boxes represent assembled pseudo-chromosomes and green boxes represent assembled scaffolds that constitute pseudo-chromosomes. **D)** BUSCO completeness scores for the sauropsida dataset and **E)** Assembly metrics (including length of the longest scaffold N50 and N90) and sequence composition (GC content) of the chromosome-level *Aldabrachelys gigantea* genome.

*Aldabrachelys gigantea* have been successfully used in rewilding projects on several Western Indian Ocean Islands, whose endemic giant tortoise species have recently gone extinct [13]. The introduced populations act as ecological replacements for the extinct species and take a central role in shaping and sustaining large-scale vegetation dynamics as the largest frugi- and herbivore[14–17]. *Aldabrachelys gigantea* have been introduced on the Mauritian islets of Ile Aux Aigrettes and Round Island and on the island of Rodrigues[18]. Monitoring the effectiveness of these rewilding projects will be crucial to catalyzing larger projects in Madagascar[19]. *Aldabrachelys gigantea* rewilding programs require genomic monitoring to reduce founder effects and maximize genetic variation in newly introduced populations[20]. Finally, uncertainties exist about the existence of additional *Aldabrachelys* lineages, as well as the number and taxonomic status of extinct lineages[10] due to weak morphological resolution and low-resolution genetic marker sets[21,22].

In this study, we present the first high-quality chromosome-level genome of *A. gigantea* using PacBio HiFi sequencing and Hi-C sequencing for scaffolding. We assessed the utility of the reference genome by performing low coverage whole-genome resequencing for a total of 32 tortoises. We inferred the genetic structure of the wild population and the likely origin of zoo-housed individuals.

## Data description

### Genome sequencing and assembly

#### DNA extraction, PacBio library preparation, and sequencing

In December 2020, during routine veterinary blood sampling, a subsample of approximately 3 mL of whole blood was collected from a female *A. gigantea* (named Hermania) living in the Zurich Zoo since 1955. Because blood was subsampled during a routine veterinary blood sampling, no additional ethical approval was required. Whole blood was taken from the animal’s dorsal tail vein and stored on ice in a heparin-coated blood collection tube. DNA extraction was carried out at the Genetic Diversity Center, ETH, Zurich, according to the manufacturer’s instructions of MagAttract® High Molecular Weight DNA (HMW) Kit (Qiagen), with a single modification: instead of using 200 µl whole blood as suggested for blood samples with non-nucleated red blood cells, a total of 50 µl whole blood was used. The purified DNA was eluted in 200 µl molecular-grade water. Subsequent steps, including gDNA quality control, PacBio HiFi library preparation, and sequencing, were carried out at the Functional Genomic Center Zurich, ETH.

The input HMW genomic DNA concentration was measured using a Qubit Fluorometer (Thermo), and the DNA integrity was checked on a Femto Pulse Device (Agilent). The HiFi library preparation started with 14 μg HMW DNA. The PacBio HiFi library was produced using the SMRTbell® Express Template Prep Kit 2.0 (Pacific Biosciences), according to the manufacturer’s instructions. Briefly, the DNA sample was mechanically sheared to an average size of 20 Kbp using a Megaruptor 3 Device (Diagenode). A Femto Pulse gDNA analysis assay (Agilent) was used to assess the resulting fragment size distribution. The sheared DNA sample was DNA damage repaired and end-repaired using polishing enzymes. PacBio sequencing adapters were ligated to the DNA template. A Blue Pippin device (Sage Science) was used to size-select fragments >15 Kbp. The size-selected library was quality inspected and quantified using a Femto Pulse gDNA analysis assay (Agilent) and a Qubit Fluorometer (Thermo), respectively. The SMRT® bell-Polymerase Complex was prepared using the Sequel® II Binding Kit 2.0 and Internal Control 1.0 (Pacific Biosciences) and sequenced on a PacBio Sequel II instrument using the Sequel II Sequencing Kit 2.0 (Pacific Biosciences). In total, two Sequel II SMRT Cells 8M (Pacific Biosciences) were run, taking one movie of 30 hours per cell. This yielded 49.4 Gbp of HiFi reads with a mean read length of 22.8 Kbp, which corresponds to approximately 20.8× coverage of the genome (Table 1).

**Table 1.** Summary of the genomic data produced in this study

***Aldabrachelys gigantea* reference genome sequencing, assembly, and validation**
|  |  |
| --- | --- |
| NCBI BioProject | PRJNA822095 |
| **Draft genome sequencing** |  |
| PacBio SMRT II HiFi data (Gb) | 192 |
| HiFi reads NCBI SRA Accession | SRR18672579 |
| **Hi-C scaffolding** |  |
| Illumina NovaSeq 6000 data (Gb) | 196 |
| Hi-C reads SRA Accession | SRR18673000 |
| **Chromosome-level genome assembly (AldGig_1.0)** |  |
| Assembled genome size (Gb) | 2.37 |
| Scaffold N50 (Mb) | 148.6 |
| No. of scaffolds | 719 |
| Contig N50 (Mb) | 61.5 |
| No. of contigs | 422 |
| BUSCO completeness (sauropsida_odb10) | 97.3% complete, 0.4% fragmented, 2.3% missing |
| **Mitochondrial genome assembly** |  |
| Assembled genome size (bp) | 16,467 |
| **Genome annotation** |  |
| PacBio SMRT IsoSeq data (Gb) | 1.1 |
| IsoSeq reads NCBI SRA Accession | SRR18674283 |
| No. of predicted protein-coding genes | 23,953 |
| No. of functionally annotated genes | 22,554 |
| BUSCO completeness (sauropsida_odb10) | 91.9% complete, 2.3% fragmented, 5.8% missing |
| **Low coverage whole-genome resequencing** |  |
| Illumina NovaSeq 6000 data (Gb) | 202 |
| NCBI SRA Accessions | SRR14611971-SRR18674101 |

#### Nuclear genome assembly, contamination scan, and evaluation

The consensus circular sequences per each Sequel II SMRT Cell (Pacific Biosciences) were filtered for adapter contamination with HiFiAdapterFilt v2.0.0 [23,24]. Overall, 0.008% of the HiFi reads were filtered out. Genome size and heterozygosity rate were estimated based on the 17-mer frequency of the cleaned HiFi reads with GCE v1.0.2 (GCE, RRID: SCR_017332) [25,26]. Our results indicate that *A. gigantea* has an estimated genome size of 2.37 Gbp (Fig. S1) and low heterozygosity of 0.072% (corresponding to 0.72 SNPs per 1 Kbp).

The reads were then assembled with the default parameters of HiCanu v2.1.1 (Canu, RRID:SCR_015880)[27,28], Improved Phased Assembler v1.3.2 (IPA HiFi Genome Assembler, RRID:SCR_021966)[29], and Hifiasm v0.15.5 (Hifiasm, RRID:SCR_021069)[30,31]. Additionally, an option for assembling inbred/homozygous genomes (-l 0) within Hifiasm (Hifiasm, RRID:SCR_021069)[30,31] was also tested. Main contiguity statistics were calculated with QUAST v5.0.2 (QUAST, RRID:SCR_001228)[32,33]. The subsequent analyses were performed with the draft assembly obtained via Hifiasm (Hifiasm, RRID:SCR_021069)[30,31] with default parameters because it provided the most contiguous and complete assembly with 483 contigs and an N50 of 61.5 Mbp (Table S1).

Scanning for contaminant contigs in the draft assembly was performed by following three approaches. First, the draft assembly was split into 5 Kbp segments using SeqKit v0.16.1 (SeqKit, RRID:SCR_018926)[34,35]. Each segment was blasted against the full NCBI non-redundant protein database by running diamond v2.0.9 (DIAMOND, RRID:SCR_016071)[36,37] with the blastx option. The decision of a segment coming from an external source was given by considering the bitscore evalue and GC content, and none of the hits were considered a significant match with cut-off values of 30, 0.0001, 70%, respectively. Second, we assessed k-mer profiles of the most probable sources of contamination human genome (NCBI RefSeq: GCF_000001405.39) and the *A. gigantea* mitochondrial genome; NCBI RefSeq: NC_028438.1), were compared with the k-mer profile of the draft assembly using KAT v2.4.1 (KAT, RRID:SCR_016741)[38,39]. The k-mer profiles were well distinct suggesting no apparent contamination. Third, the previously published *A. gigantea* whole-genome resequencing dataset (NCBI SRA: SRX4741543)[12] was mapped against our assembly with BWA-MEM v0.7.17 (BWA, RRID:SCR_010910)[40]. The read coverage profile was examined with Qualimap v2.2.1 (QualiMap, RRID:SCR_001209)[41,42]. The resequencing dataset had 27× coverage, therefore we discarded contigs from the assembly with less than 10× or more than 100× aligned read depth. All contaminant filtering steps combined, 62 contigs were removed from the assembly, resulting in a final set of 422 contigs and an N50 of 61.5 Mbp (Table S1).

We assessed the completeness of the assembly based on a BUSCO analysis of single-copy orthologs v5.1.2 (BUSCO, RRID:SCR_015008)[43,44] with default parameters and the sauropsid dataset (sauropsida_odb10) in the genome mode.

#### Hi-C sequencing and genome scaffolding

The Hi-C library was constructed with a 250 µl whole blood sample that was first fixed with 1% formaldehyde for 15 min at room temperature. Then, solid glycine powder was added to obtain a final concentration of 125 mM and incubated for 15 min at room temperature with periodic mixing. After centrifugation, the pellet was resuspended in PBS + 1% Triton-X solution and incubated at room temperature for 15 min. Then, the nuclei were collected after the mixture was spun down. The cross-linked sample was sent on dry ice to Phase Genomics (Seattle, Washington, USA) for sequencing. The Hi-C library was generated using the Phase Genomics Proximo Animal kit version 4.0. Briefly, the DNA sample was digested with DpnII and the 5’-overhangs were filled while incorporating a biotinylated nucleotide. The blunt-end fragments were ligated, sheared, and the biotinylated ligation junctions captured with streptavidin beads. The resulting fragments were sequenced on a NovaSeq 6000 150 bp paired-end run. A total of 680 million reads were produced, corresponding to approximately 85× coverage of the genome (Table 1).

Overall, 90.3% of the Hi-C reads were aligned to the draft genome assembly, sorted, and merged. Then duplicates were removed using Juicer v1.6 (Juicer, RRID:SCR_017226)[45,46] with default parameters. Approximately 87% of the reads were found to have Hi-C contacts. Afterward, the 3D-DNA pipeline was run with default parameters to generate a candidate assembly[47,48], which was reviewed using JBAT v2.10.01[49]. Finally, a high-quality chromosome-level genome assembly was generated after a visual review on JBAT[49]. A total of 26 pseudo-chromosomes were anchored, corresponding to 97.6% of the estimated genome size, yielding a chromosome-level assembled reference genome with an N50 of 148.6 Mbp (Table 1, Fig. 1C) and a BUSCO completeness of 97.3% (Fig. 1D, Table 1). Genome assembly statistics were visualized with a snail plot in BlobToolKit v2.6.4 [50,51] (Fig. 1E). The chromosome level assembly of *A. gigantea* (AldGig_1.0) has the longest contig and scaffold N50, and one of the highest BUSCO completeness scores of all available chromosome-level assembled chelonian genomes (Table 2).

**Table 2.** Contiguity and completeness statistics of all available chromosome-level assembled chelonian genomes

| Species name<br>(Accession No) | Family | Genome size<br>(Gbp) | Contig N50<br>(Mbp) | Scaffold N50<br>(Mbp) | BUSCO completeness* |
| --- | --- | --- | --- | --- | --- |
| *Aldabrachelys gigantea*<br>(This study) | Testudinidae | 2.374 | 61.5 | 148.6 | 97.3%[S:96.2%,D:1.1%],F:0.4%,M:2.3% |
| *Gopherus evgoodei*<br>(GCF_007399415.2) | Testudinidae | 2.299 | 13.027 | 147.4 | 97%[S:95.9%,D:1.1%],F:0.5%,M:2.5% |
| *Chelonia mydas*<br>(GCF_015237465.2) | Cheloniidae | 2.134 | 39.416 | 134.4 | 97.3%[S:96.3%,D:1.0%],F:0.4%,M:2.3% |
| *Dermochelys coriacea*<br>(GCF_009764565.3) | Dermochelyidae | 2.165 | 7.03 | 137.6 | 96.3%[S:95.3%,D:1.0%],F:0.6%,M:3.1% |
| *Chrysemys picta bellii*<br>(GCF_000241765.4) | Emydidae | 2.481 | 0.021 | 16 | 96.6%[S:95.7%,D:0.9%],F:1.1%,M:2.3% |
| *Trachemys scripta elegans*<br>(GCF_013100865.1) | Emydidae | 2.126 | 0.205 | 140.4 | 95.0%[S:94.0%,D:1.0%],F:1.2%,M:3.8% |
| *Mauremys mutica*<br>(GCF_020497125.1) | Geoemydidae | 2.484 | 15.011 | 135 | 97.3%[S:95.2%,D:2.1%],F:0.5%,M:2.2% |
| *Mauremys reevesii*<br>(GCF_016161935.1) | Geoemydidae | 2.368 | 33.353 | 130.5 | 97.5%[S:95.9%,D:1.6%],F:0.4%,M:2.1% |
| *Rafetus swinhoei*<br>(GCA_019425775.1) | Trionychidae | 2.238 | 30.964 | 132 | 96.4%[S:95.3%,D:1.1%],F:0.6%,M:3.0% |

\*BUSCO score generated from the sauropsid (sauropsida\_odb10) database. BUSCO statistics C, complete; S, single-copy; D, duplicated; F, fragmented; M, missing

#### Repetitive element analysis

To identify, classify, and mask repetitive elements in the *A. gigantea* genome, we first generated a species-specific *de novo* repeat library using RepeatModeler v2.0.1 (RepeatModeler, RRID:SCR_015027)[52,53]. RepeatModeler utilizes RECON (RECON,RRID:SCR_021170)[54], RepeatScout (RepeatScout, RRID:SCR_014653)[55], and Tandem Repeats Finder (Tandem Repeats Database, RRID:SCR_005659)[56] to detect repeat families *de novo*, to identify and classify consensus sequences. These consensus sequences were then used to softmask the genome with RepeatMasker v4.1.0 (RepeatMasker, RRID:SCR_012954)[57]. As a result, 46.7% of the genome (1,114,704,617 bp) were detected as repetitive and softmasked. Long interspersed nuclear elements (LINEs) were identified as the most abundant class of repetitive elements (12.36%) followed by long terminal repeat (LTR) elements (5.78%) (Table S2).

#### RNA extraction and sequencing

A whole blood sample of approximately 1 mL was collected from an individual named Grosser Bub (‘Big Boy’) during routine veterinary blood sampling in the Zurich Zoo. A total of 125 µl of whole blood was immediately diluted with the same amount of water, added into TRIzol™ LS Reagent (Invitrogen), and stored on ice for < 2 hours until extraction. RNA was extracted at the Genetic Diversity Center, ETH, following a combination of a TRIzol™ LS (Invitrogen) RNA isolation protocol and the RNeasy Mini Kit (Qiagen). First, the sample was incubated at room temperature for 5 min. Then, 0.2 mL chloroform was added to the sample and the mixture was inverted for 15 seconds, followed by 3 min incubation at room temperature. The resulting mixture was centrifuged at 11,000 rpm for 15 min at 4°C. After centrifugation, the upper phase containing the RNA was collected, mixed with 1x 70% ethanol, and transferred to an RNeasy spin column. For the remaining procedure, the protocol “Purification of Total RNA from Animal Tissues” of the kit was followed, starting from step 6. Briefly, the RNA was bound to the spin column, washed, and eluted in 30 μl molecular grade water. The initial quality control of the RNA was done on a TapeStation (Agilent) and the concentration was measured with a Qubit Fluorometer (Thermo).

The PacBio IsoSeq library for RNA sequencing was produced at the Functional Genomic Center Zurich using the SMRTbell Express Template Prep Kit 2.0 (Pacific Biosciences), according to the manufacturer’s instructions. A total of 300 ng RNA was used as input for the cDNA synthesis, which was carried out using the NEBNext® Single Cell/Low Input cDNA Synthesis & Amplification Module (NEB) and Iso-Seq Express Oligo Kit (Pacific Biosciences) following instructions. To enrich for longer transcripts (>3 kb), 82 µl ProNex Beads were used for the clean-up of the amplified DNA, as outlined in the protocol. For all subsequent quality control steps, a Bioanalyzer 2100 12 Kb DNA Chip assay (Agilent) and a Qubit Fluorometer

(Thermo) were used to assess the size and concentration of the library. The SMRT bell-Polymerase Complex was prepared using the Sequel Binding Kit 3.0 (Pacific Biosciences) and sequenced on a PacBio Sequel instrument using the Sequel Sequencing Kit 3.0 (Pacific Biosciences). In total, one Sequel™ SMRT® Cell 1M v3 (Pacific Biosciences) was run with one movie of 20 hours per cell producing ∼1.1 Gbp of HiFi data (Table 1).

#### Gene prediction and annotation

Gene prediction was performed using a combination of *ab initio* and evidence-based prediction methods (RNA-seq and homology-based) with the braker2 pipeline v2.1.5 (BRAKER, RRID:SCR_018964)[58–62]. All gene predictions were performed with pre-trained parameter sets for chicken, which is the evolutionarily closest taxon for *A. gigantea* available within the software. Using pre-trained parameters yielded more complete annotations compared to training with extrinsic evidence (i.e. RNA-seq and protein data). The *ab initio* prediction was performed by utilizing the soft-masked reference genome. Evidence for the transcriptome-based prediction was based on combining information from *A. gigantea* PacBio Iso-seq and all available RNA-seq databases from chelonians in closely related genera (*Chelonoidis* spp. and *Gopherus* spp.; Table S3). For the alignment of short and long read transcripts, the splice-aware alignment tools STAR v2.7.9 (STAR, RRID:SCR_004463)[63,64] and minimap2 v2.24 (Minimap2, RRID:SCR_018550)[65,66] were used, respectively. Additionally, evidence for the homology-based prediction consisted of a protein database combining all vertebrate proteins in the OrthoDB v10 (OrthoDB, RRID:SCR_011980)[67] and the protein sequences of *Gopherus evgoodei* (NCBI RefSeq: GCF_007399415.2) and *Chelonoidis niger abingdonii* (NCBI RefSeq: GCF_003597395.1). This dataset was aligned against the chromosome-level assembled reference genome via the ProtHint pipeline v2.6.0 (ProtHint, RRID:SCR_021167)[68,69]. RNA-seq and homology-based evidence were incorporated for the braker2 pipeline (BRAKER, RRID:SCR_018964) run in --etpmode[62,68,70–74]. All gene models derived from *ab initio* and evidence-based methods were integrated into a high confidence non-redundant gene set by using TSEBRA v1.0.3 [75,76], with the “keep ab_initio” configuration set. The translated protein sequences from the predicted gene models were searched against protein profiles corresponding to major clades/families of transposon open reading frames by TransposonPSI v1.0 [77]. Overall, 331 genes were identified as likely derived from transposable elements and excluded from the annotation. The resulting gene model set consisted of 23,953 protein-coding genes with a mean gene length of 39,458 bp and an average of nine exons per coding sequence (Table 1).

The completeness of the annotation was assessed based on single-copy orthologs via BUSCO v5.1.2(BUSCO, RRID:SCR_015008)[43,44] with default parameters in the protein mode. The proteome BUSCO completeness scores were 93.7% and 91.9% for the vertebrate (vertebrata_odb10) and sauropsida (sauropsida_odb10) datasets, respectively. The level of BUSCO completeness for the datasets is comparable to those of the annotations of the *C. n. abingdonii* (vertebrata, 96.9%; sauropsida, 97.7%) and *G. evgoodei* (vertebrata, 99.7%; sauropsida, 99.3%) (Table S4).

Functional annotation of the encoded proteins was performed using the suite of search tools included in InterProScan v5.53-87.0 (InterProScan, RRID:SCR_005829)[78,79], with default parameters, in combination with putative gene names derived from UniProtKB/Swiss-prot (UniProtKB/Swiss-Prot, RRID:SCR_021164)[80]. AGAT v0.8.0[81,82] was used for summarizing the properties of the structural annotation and for combining the structural and functional annotation results. Of all the prediction gene models, 94.1% could be functionally annotated (Table 1, Table S5).

#### Identification of non-coding RNA genes

tRNA, rRNA, snRNA, and miRNA were annotated using Infernal v1.1.4 (Infernal, RRID:SCR_011809)[83,84], which builds covariance models as consensus RNA secondary structure profiles from the genome. The tool then uses the models to search Rfam (Rfam, RRID:SCR_007891)[85], a database of non-coding RNA families. Overall, the homology-based non-coding RNA annotation revealed a total of 6,754 tRNAs, 3,636 rRNAs, 345 snRNAs, and 671 miRNAs encoded in the genome.

#### Mitochondrial genome assembly and evaluation

Mitochondrial reads were extracted from the PacBio HiFi dataset and assembled with Hifiasm v0.15.5 (Hifiasm, RRID:SCR_021069)[30,31] using the MitoHifi v2.0 pipeline[86,87]. The genome size was 16,467 bp and the assembly was 100% identical at the nucleotide level to the *A. gigantea* mitochondrial reference genome available at NCBI RefSeq with accession no NC_028438.1[88].

### Synteny analysis

We investigated the collinearity of *A. gigantea* chromosomes with three other chromosome-level chelonian genome assemblies from three different families including *Gopherus evgoodei* (Testudinidae; NCBI RefSeq: GCF_007399415.2), *Mauremys mutica* (Geoemydidae; NCBI RefSeq: GCF_020497125.1) and *Trachemys scripta elegans* (Emydidae; NCBI RefSeq: GCF_013100865.1) (Fig. 2). We analyzed the largest 10 chromosomes corresponding to 75% of the *A. gigantea* assembly. Chromosomes from each genome were aligned to other genomes using minimap2 v2.24 (Minimap2, RRID:SCR_018550)[65,66] with default parameters (-ax asm5). The resulting alignments were processed with SyRI v1.5.4 [89,90] to identify syntenic regions and structural rearrangements. The syntenic regions and structural rearrangements for the four chelonian genomes were visualized with plotsr v0.5.3[91,92]. Among genomes, we found between 1.5–1.6 Gbp of syntenic regions and 15.3–54.6 Mbp of rearrangements corresponding to 89–94% and 0.8–3% of the compared genome portions, respectively. The rearrangements included 0.2–1.4 Mbp of duplications, 2.5–7.7 Mbp of translocations, and 6.6– 51.6 Mbp of inversions. The high ratio of syntenic regions that we found are between chelonian taxa that diverged around 50–70 mya[93] (Fig. 2) and is in agreement with previous studies, where the base substitution rate (evolutionary rate) of chelonians was found to be relatively low (see[94]).

**Fig. 2.**
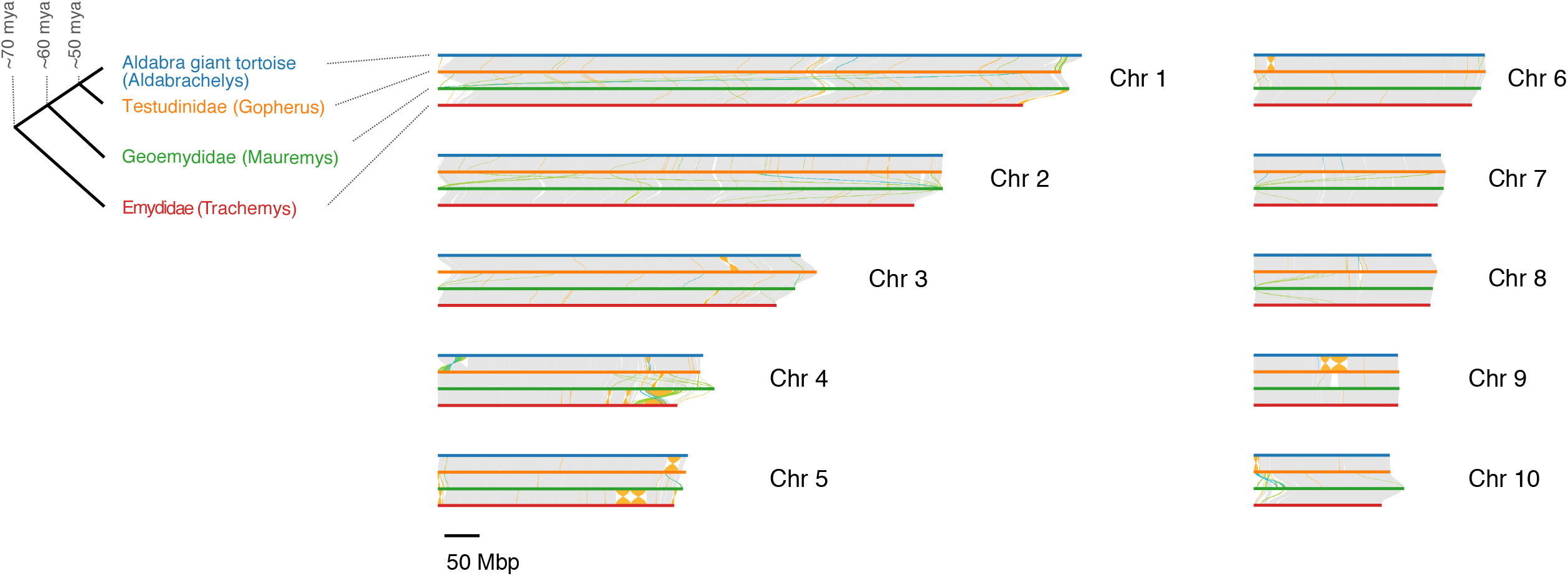
Synteny analysis of 10 chromosomes in *Aldabrachelys gigantea* (blue horizontal lines), *Gopherus evgoodei* (orange horizontal lines), *Mauremys mutica* (green horizontal lines), and *Trachemys scripta elegans* (red horizontal lines) genome assemblies shows high levels of conservation between distantly related chelonian taxa. Gray, yellow, green, and light blue lines between assemblies indicate syntenic regions, inversions, translocations, and duplications, respectively. The four compared assemblies represent all chelonian families (see cladogram on the left, split times from [93]) except Platysternidae within the chelonian super family Testudinoidea, which includes families of Emydidae (terrapins), Geoemydidae and Testudinidae (land-dwelling tortoises). *Trachemys scripta elegans* is from Emydidae, *Mauremys mutica* is from Geoemydidae, and *Gopherus evgoodei* and *Aldabrachelys gigantea* are from Testudinidae.

We also performed a complementary collinearity analysis based on orthologous gene sets of *A. gigantea* and the phylogenetically closest available chromosome-level assembled *Gopherus evgoodei* (NCBI RefSeq: GCF_007399415.2) reference genomes (split time ca. 50 my[93]). We first created orthogroups with the proteomes of the two species using Orthofinder v2.5.4 (OrthoFinder, RRID:SCR_017118)[95,96]. A total of 41979 genes (91.3% of total) were assigned to 15662 orthogroups. The orthologues were then fed in i-ADHoRe v.3.0 [97,98] to detect genomic regions with statistically significant conserved gene content requiring a minimum of three anchor points within each syntenic region (gap_size=15, cluster_gap=30, q_value=0.05, prob_cutoff=0.01, anchor_points=3, alignment_method=gg2, level_2_only=true). Finally, longer-term ancestral synteny detected for the two species was visualized with Circos v0.69-8 (Circos, RRID:SCR_011798)[99,100](Fig. S2). Eventually, the high ratio of synteny revealed by our complementary collinnearity analysis supported our findings on the chromosomal synteny analysis.

### Sample collection for low coverage whole-genome resequencing

The native distribution of *A. gigantea* is restricted to the Aldabra Atoll (Fig. 1B) with deep water channels separating the four main islands (Grande Terre, Malabar, Polymnie, and Picard; Fig. 3A). The smallest island Polymnie no longer harbors any tortoises[101]. Tortoises were also harvested to extinction on Picard in the 1800s but the island has since been re-populated through translocations from Malabar and Grande Terre[102]. In addition to the Atoll, there is an unknown, but large number of ex-situ individuals in captive, semi-natural, or rewilded populations [13]. Assessments of the genetic health of native and rewilded populations will be crucial to inform future species management. However, the uncertainty about genomic vulnerabilities and which ex-situ individuals to use for re-wilding efforts constitute significant barriers.

**Fig. 3.**
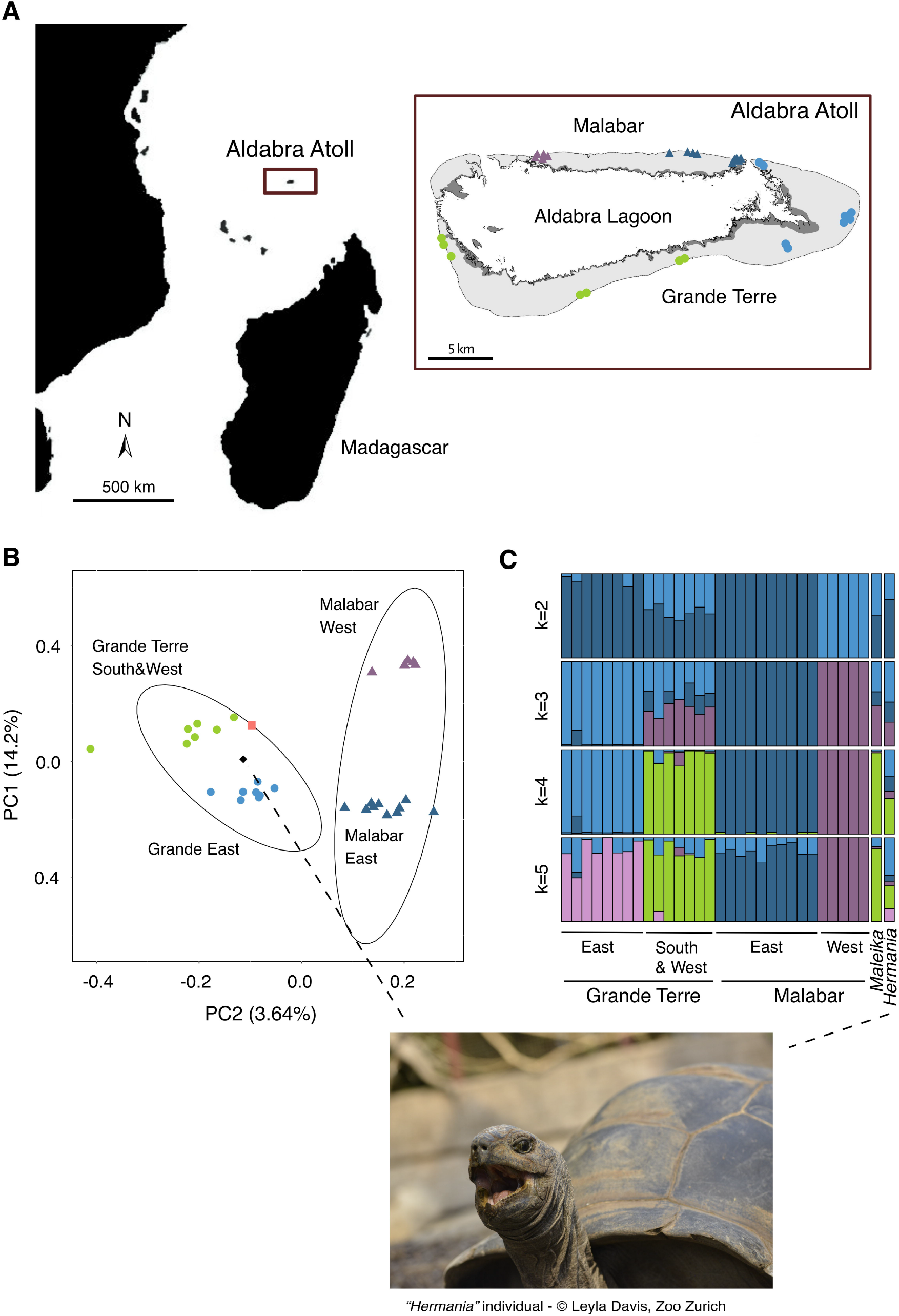
**A)** The location of the Aldabra Atoll in the Western Indian Ocean and sampling location of 30 individuals within Aldabra Atoll. Every colored mark on the map represents a sampled tortoise. **B)** Principal component analysis plot of 30 *Aldabrachelys gigantea* individuals from the islands of Grande Terre (East, light blue circles; South & West, green circles) and Malabar (East, dark blue triangles; West, purple triangles) and two individuals from the Zurich Zoo (Hermania, black diamond; Maleika, light pink square). Principal components 1 and 2 account for 14.2% and 3.64% of the overall genetic variability, respectively. **C)** Admixture proportions of all the individuals for ancestral populations (k) varied from 2 to 5. Each bar represents one individual and shows its admixture proportions.

To assess the utility of our reference genome resources to improve the genomic monitoring and inform re-wilding efforts, we performed low-coverage whole-genome sequencing of a representative sample of two main islands as well as zoo-housed individuals. Low-coverage sequencing is a powerful and cost-effective approach for conservation and population genomics [103], as well as ancient DNA analyses [104]. We collected blood samples from a total of 30 adult *A. gigantea* (Table S6) from Malabar (East, N=10; West, N=5) and Grande Terre (East, N=8; South, N=4; West, N=3) (Fig. 3A). The collection yielded ∼200 µl of blood from the cephalic vein of a front limb. We received a research permit from the Seychelles Bureau of Standards (ref #A0347) for our collection. European institutions currently host over 360 *A. gigantea* [105]. Here, we analyzed two female individuals living in Zurich Zoo, Switzerland. The individual named Hermania was used to create the reference genome, and the individual named Maleika arrived at Zurich Zoo in 1984 and lived there until her death in 2018. The historic information surrounding the exact importation location from Aldabra is sparse or unknown. Sampling from Hermania was performed as described above, and sampling from Maleika was performed by using ∼500 mg of muscle tissue sampled after veterinary necropsy and stored in absolute ethanol until DNA extraction.

### DNA extraction, and sequencing

DNA extraction was performed with 3 µl of blood from Hermania and 15 mg of muscle tissue from Maleika, using the sbeadex™ kit (LGC Genomics, Middlesex, UK), following the manufacturer’s protocol for DNA extraction from nucleated red blood cells and tissue, respectively. Genomic DNA concentrations were measured with a dsDNA Broad Range Assay Kit (Qubit 2.0 Fluorometer, Invitrogen, Carlsbad). More than 200 ng of DNA per sample was sent to Novogene Company (Cambridge, United Kingdom) for library preparation and sequencing. Briefly, the genomic DNA was randomly fragmented to a size of 350 bp, end-polished, A-tailed and ligated with Illumina adapters of Illumina sequencing. After PCR enrichment, products were purified (AMPure XP system) and checked for quality on an Agilent 2100 Bioanalyzer (Agilent Technologies, CA, USA). Molarity was assessed using real-time PCR. Libraries were sequenced on the Illumina Novaseq 6000 platform with paired-end runs of 150 bp read length. For each of the 32 samples, ∼2.6 Gbp raw reads were generated (Table S6, Table 1).

### Data filtering, alignment, and genotype likelihood estimation

To account for the low coverage sequencing approach, we assessed genotype likelihoods using the Atlas Pipeline [106,107]. We first used the GAIA workflow to remove Illumina adapters with TrimGalore v0.6.6 (Trim Galore, RRID:SCR_011847)[108] with default parameters. Only reads longer than 30 bp were retained. Then, reads were aligned to the reference genome with BWA using BWA-MEM v0.7.17 (BWA, RRID:SCR_010910)[40] filtering for mapping quality scores >20. Alignments were processed with the RHEA workflow for indel realignment with GATK v3.8 (GATK, RRID:SCR_001876)[109]. A target interval set was created with a representative set of 15 samples and each individual was realigned together with a representative set of individuals (guidance samples) to enable realignment of low coverage samples without jointly realigning all samples. The average read depth per sample was 1.62– 2.06 with a mean of 1.79 (Table S6).

We used ANGSD v0.93 (ANGSD, RRID:SCR_021865)[110,111] to produce genotype likelihoods appropriate for the low coverage of individual samples. GATK was used (GATK, RRID:SCR_001876)[109] to infer major and minor alleles from the likelihoods (doMajorMinor 1, doMaf 1). Quality filtering for the subsequent downstream analyses was performed as follows: Only properly paired (only_proper_pairs 1) and unique reads (uniquieOnly 1) were used, and only biallelic sites were retained (skipTrialleleic 1). Nucleotides with base qualities below 20 were discarded. Excessive SNPs around indels and excessive mismatches with the reference were corrected (C50, baq 1,[112]. Sites with read coverage in fewer than 50% of the samples were excluded (minimum representation among samples > 50%, -minInd 16). SNPs with a genotype likelihood p-value < 0.001 were retained, producing a final set of 7,131,506 variant sites.

### Population genetic structure and individual assignments

Our low coverage sequencing analyses focused on revealing within and among island genetic differentiation in the Aldabra Atoll, as well as assigning likely origins for zoo-housed individuals. We first assessed the global genetic structure of the samples using a principal component analysis with PCAngsd v09.85[113]. Based on a total of 6,651,907 variant sites with a minor allele frequency >0.05, individuals from Malabar and Grande Terre split into individual groups (Fig. 3B). Both zoo samples were grouped within the group of Grande Terre individuals revealing the most likely origin for these individuals captured in the 20th century. The principal component analysis also reveals a finer scale east-west population structure within islands confirming recent results based on ddRAD sequencing [114].

We also assessed genetic structure using unsupervised Bayesian clustering with NGSAdmix (NGSadmix, RRID:SCR_003208)[115]. We performed pairwise linkage disequilibria (LD) pruning to reduce dependence among SNP loci[115]. Pairwise LD was calculated using ngsLD[116] and LD pruning was performed by allowing a maximum among-SNP distance of 100 Kbp and a minimum weight of 0.5. After LD pruning, 5,862,629 SNPs were retained and 50 replicate runs of NGSAdmix (NGSadmix, RRID:SCR_003208)[115] were performed. We varied the number of clusters (k) between 2-5 and visualized the assignments with PopHelper v.1.0.10[117,118] (Fig. 3C). The admixture analyses for k=2 clusters revealed a main split with groups formed by East Grande Terre together with East Malabar opposed to West Malabar. South & West Grande Terre individuals were assigned to both groups. At k=4 each major sampling region was assigned to a single cluster. The zoo individual Maleika showed a genotype highly consistent with South & West Grande Terre individuals. The individual Hermania was assigned to different Grande Terre regions.

## Conclusions

We assembled the first high-quality, chromosome-level annotated genome for the Aldabra giant tortoise, resulting in one of the best assembled chelonian genomes. Chromosomal collinearity analyses revealed a high degree of conservation even among distantly related tortoise species. We showed that the high-quality resources can be combined with low-coverage resequencing to gain crucial insights into the genetic structure within the Aldabra Atoll, as well as to resolve the exact origin of zoo-housed individuals. Understanding levels of genomic diversity in both native and ex-situ populations is crucial to inform rewilding efforts and prioritize conservation efforts. Furthermore, genome-wide analyses of polymorphism can be used to assess the presence of deleterious mutations endangering the long-term health of populations. Finally, given the exceptionally long lifespan and large body size, the high-quality genome will inform comparative genomics studies focused on the genetic underpinnings of aging and gigantism.

## Data availability

The raw sequencing data and the nuclear and mitochondrial genome assemblies produced in this study have been deposited in the NCBI under BioProject accession number PRJNA822095.

## Abbreviations

μg: microgram
μl: microliter
°C: degree Celcius
AGAT: Another Gtf/Gff Analysis Toolkit
ANGSD: Analysis of Next Generation Sequencing Data
baq: base alignment quality
bp: base pairs
BUSCO: Benchmarking Universal Single-Copy Orthologs
BWA: Burrows-Wheeler Aligner
DNA: deoxyribonucleic acid
cDNA: complementary DNA
dsDNA: double-strand DNA
EAZA: European Association of Zoos and Aquaria
ETH: Swiss Federal Institute of Technology in ZüCrich
GAIA: Genome-wide Alignment Including Adapter-trimming
GATK: Genome Analysis Toolkit
Gbp: gigabase pairs
GC: guanine and cytosine
GCE: Genomic Character Estimator
gDNA: genomic DNA
Hi-C: chromosome conformation capture
HiFi: high-fidelity
HMW: high molecular weight
IsoSeq: isoform sequencing
IUCN: International Union for Conservation of Nature
JBAT: Juicebox Assembly Tools
KAT: k-mer analysis toolkit
LS: liquid sample
MAF: minor allele frequency
Mbp: megabase pairs
mg: milligram
min: minute
mL: milliliter
mM: millimolar
NCBI: National Center for Biotechnology Information
NEB: New England Biolabs
ng: nanogram
NGSAdmix: Next Generation Sequencing Admixture
ngsLD: Next Generation Sequencing Linkage Disequilibrium
OrthoDB: orthologous database
PacBio: Pacific Biosciences
PBS: phosphate-buffered saline
PCA: principal component analysis
PCR: polymerase chain reaction
QUAST: Quality Assessment Tool
RefSeq: reference sequence
Rfam: RNA families
RNA: ribonucleic acid
RNA-seq: RNA sequencinf
rpm: revolutions per minute
Sauropsida_odb10: sauropsids orthologous database 10
SMRT: single-molecule real-time
SNP: single nucleotide polymorphism
SRA: Sequence Read Archive
STAR: Spliced Transcripts Alignment to a Reference
SyRI: Synteny and Rearrangement Identifier
tRNA: transfer RNA
rRNA: ribosomal RNA
snRNA: small nuclear RNA
miRNA: micro RNA
TSEBRA: Transcript Selector for BRAKER
UniProtKB: Universal Protein Knowledgebase
Vertebrata_odb10: vertebrate orthologous database 10

## Competing interests

The authors declare that they have no competing interests.

## Funding

This study was funded through the Research Talent Development Fund of the University of Zürich, the Swiss National Science Foundation (Project No: 31003A_182343), and University of Zurich Internal Funds, all of which were given to C.G.

## Author contributions

F.G.*Ç*. and C.G conceived the study design. F.G.*Ç*. carried out all DNA and RNA extractions and bioinformatic analyses with guidance from D.C. and C.G. D.H. and L.D. coordinated the sampling of the zoo animals. N.B. helped get permits required for tortoise sampling on Aldabra Atoll and L.A. managed the collection, storage, and transport of samples from wild individuals. F.G.*Ç*. wrote the manuscript with guidance from C.G and substantial input from D.C. All authors revised the manuscript.

## Acknowledgments

We gratefully acknowledge Jean-Michel Hatt, Gabriela Hurlimann, and Maya Kummrow for facilitating sample donation, Claudia Rudolf von Rohr, and the team of tortoise keepers at the Masoala Rainforest in Zurich Zoo for their assistance and helpful discussions. We thank the Seychelles Islands Foundation staff Maria Bielsa, Mickael Esparon, Bruno Mels, Martin van Rooyen, Mersiah Rose, and Brian Souyana for sample collection on Aldabra Atoll; Ronny Rose and Frauke Fleischer-Dogley for their assistance with the handling, storage, and transport of the samples. We also thank Sirpa Kurz from the Zoological Museum, University of Zurich, and Constantin Latt from the Natural History Museum of Bern for their help in tissue sampling of the tortoise Maleika. Additionally, we thank Silvia Kobel and Aria Minder from the Genetic Diversity Centre, ETH Zurich for their help in wet lab applications. We also thank Simon Grüter and Weihong Qi from the Functional Genomics Center, ETH, Zurich for their help in getting sequencing services.

## Notes

### Competing Interest Statement

The authors have declared no competing interest.

