## Supplementary material for "Chromosome-level genome assembly for the Aldabra giant tortoise enables insights into the genetic health of a threatened population": Fig. S1, Fig. S2

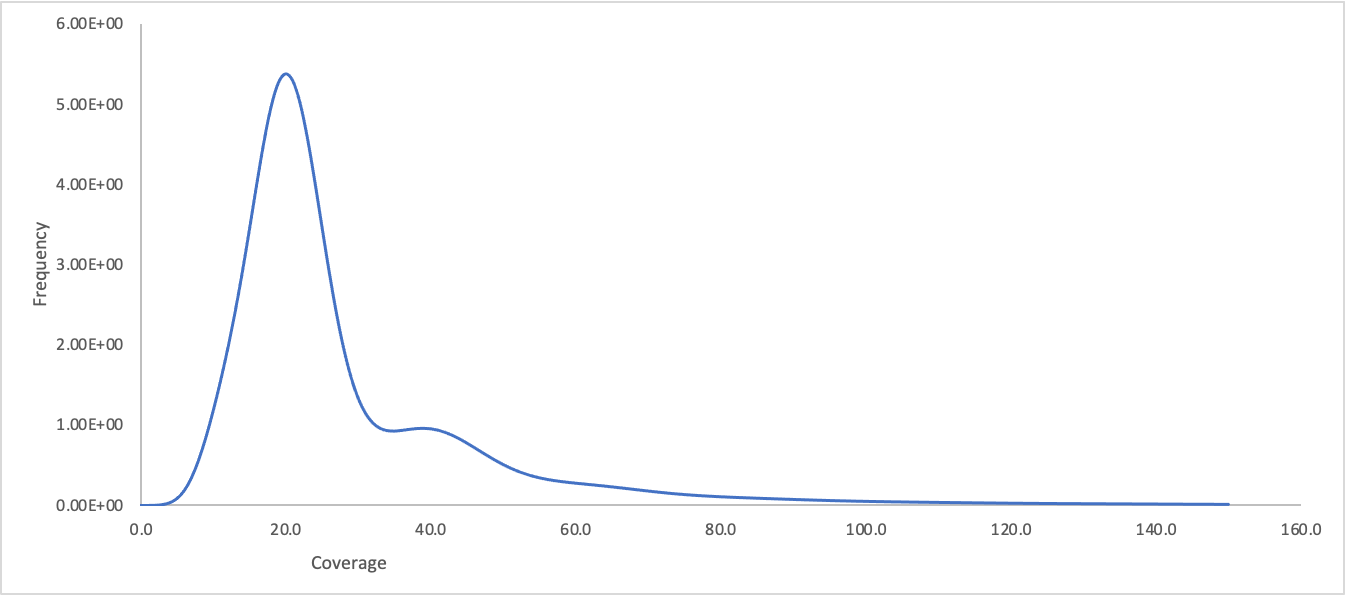

**Fig. S1** K-mer (k=17) profile of the *Aldabrachelys gigantea* genome. Consistent with low heterozygosity, most of the k-mers form one peak centered around roughly 20× coverage and do not form another peak centered at roughly half the coverage that would represent k-mers arising from heterozygous alleles.

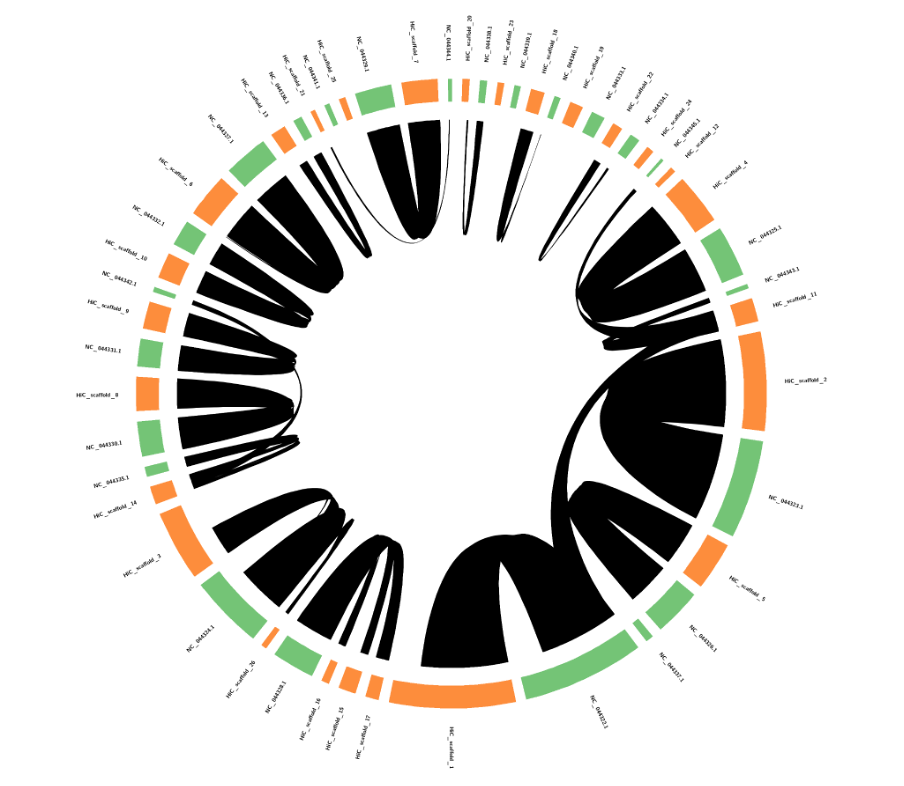

**Fig. S2** Circos plot showing the synteny between the *Aldabrachelys gigantea* Hi-C scaffolds (orange) and *Gopherus evgoodei* assembly pseudo-chromosomes (green).
