## Supplementary material for "Chromosome-level genome assembly for the Aldabra giant tortoise enables insights into the genetic health of a threatened population": Table S1, Table S2, Table S3, Table S4, Table S5

**Table S1** Genome contiguity statistics of the assemblies obtained from different assemblers. The column shaded in gray represents our initial assembly obtained via default parameters in Hifiasm

| **Genome Statistics** | **HiCanu** | **IPA** | **Hifiasm**  **(default)** | **Hifiasm**  **(with -l 0 option)** |
| --- | --- | --- | --- | --- |
| Contig n | 867 | 2446 | 422 | 703 |
| Contig N50 | 12.6Mbp | 1.6Mbp | 61.5Mbp | 41.4Mbp |
| N Count | 0 | 897 | 0 | 0 |
| Largest contig | 49.8Mbp | 7.8Mbp | 210.3Mbp | 140.8Mbp |
| Total length | 2.3Gbp | 2.3Gbp | 2.4Gbp | 2.5Gbp |
| GC (%) | 44.2 | 44.0 | 44.4 | 44.5 |

**Table S2** Summary of repeat annotations

| **Type** | **Length (bp)** | **% in genome** | **# of elements** |
| --- | --- | --- | --- |
| Retroelements | 482,092,777 | 20.31 | 1,177,209 |
| LINEs | 293,395,900 | 12.36 | 695,701 |
| LTRs | 137,235,010 | 5.78 | 154,762 |
| SINEs | 51,461,867 | 2.17 | 326,746 |
| DNA elements | 198,183,931 | 8.35 | 642,321 |
| Unclassified | 407,271,311 | 17.16 | 1,902,917 |
| Total interspersed repeats | 1,087,548,019 | 45.82 |  |

**Table S3** Accession details of the short read RNA-seq samples used in this study

| **BioSample ID** | **Sample Type** | **Species** | **Reference** |
| --- | --- | --- | --- |
| SAMN03496275 | Whole blood | *Gopherus evgoodei* | Rhie et al. 2021  Koepfli et al. 2015 |
| SAMN03496276 | Whole blood | *Gopherus morafkai* | NA |
| SAMN03496277 | Whole blood | *Gopherus agassizii* | NA |
| SAMN03496278 | Whole blood | *Gopherus morafkai* | NA |
| SAMN03496279 | Whole blood | *Gopherus morafkai* | NA |
| SAMN03496280 | Whole blood | *Gopherus evgoodei* | NA |
| SAMN03496281 | Whole blood | *Gopherus evgoodei* | NA |
| SAMN03496282 | Whole blood | *Gopherus agassizii* | NA |
| SAMN03496283 | Whole blood | *Gopherus agassizii* | NA |
| SAMN02422540 | Generic sample | *Chelonoidis niger* | NA |
| SAMN02422541 | Generic sample | *Chelonoidis niger* | NA |
| SAMN02422542 | Generic sample | *Chelonoidis niger* | NA |
| SAMN02422543 | Generic sample | *Chelonoidis niger* | NA |
| SAMN02422544 | Generic sample | *Chelonoidis niger* | NA |
| SAMN02422545 | Generic sample | *Chelonoidis carbonarius* | NA |
| SAMN02800025 | Whole blood | *Chelonoidis niger* | Whitelaw et al. 2016  Romiguier et al. 2014 |
| SAMN02800026 | Whole blood | *Chelonoidis niger* | Whitelaw et al. 2016  Romiguier et al. 2014 |
| SAMN02800027 | Whole blood | *Chelonoidis niger* | Whitelaw et al. 2016  Romiguier et al. 2014 |
| SAMN02800028 | Whole blood | *Chelonoidis niger* | Whitelaw et al. 2016  Romiguier et al. 2014 |
| SAMN02800029 | Whole blood | *Chelonoidis niger* | Whitelaw et al. 2016  Romiguier et al. 2014 |
| SAMN05991310 | Skeletal muscle | *Gopherus agassizii* | NA |
| SAMN05991317 | Lung | *Gopherus agassizii* | NA |
| SAMN05991318 | Brain | *Gopherus agassizii* | NA |
| SAMN07840320 | Whole blood | *Chelonoidis abingdonii* | Quesada et al. 2019 |
| NA | Granuloma | *Aldabrachelys gigantea* | Quesada et al. 2019 |

**Table S4** BUSCO statistics for the protein coding gene annotation of *Aldabrachelys gigantea*, *Chelonoidis abingdonii*, and *Gopherus evgoodei*

| **Species**  **(GenBank Accession No)** |  | | | | |
| --- | --- | --- | --- | --- | --- |
|  |  | **Vertebrate***  **(n = 3354)** | | **Sauropsida***  **(n = 7480)** | |
|  |  | **%** | **#** | **%** | **#** |
| *Aldabrachelys gigantea* | **C** | 93.7 | 3144 | 91.9 | 6874 |
|  | **S** | 90.8 | 3046 | 88.4 | 6613 |
|  | **D** | 2.9 | 98 | 3.5 | 261 |
|  | **F** | 3.1 | 105 | 2.3 | 172 |
|  | **M** | 3.2 | 105 | 5.8 | 434 |
| *Chelonoidis abingdonii*  (GCF_003597395.1) | **C** | 96.9 | 3250 | 97.7 | 7308 |
|  | **S** | 46.1 | 1545 | 46.4 | 3470 |
|  | **D** | 50.8 | 1705 | 51.3 | 3838 |
|  | **F** | 2.5 | 83 | 1.1 | 79 |
|  | **M** | 0.6 | 21 | 1.2 | 93 |
| *Gopherus evgoodei*  (GCF_007399415.2) | **C** | 99.7 | 3342 | 99.3 | 7427 |
|  | **S** | 43.1 | 1445 | 43.8 | 3278 |
|  | **D** | 56.6 | 1897 | 55.5 | 4149 |
|  | **F** | 0.1 | 4 | 0.1 | 10 |
|  | **M** | 0.2 | 8 | 0.6 | 43 |

*BUSCO score generated from the vertebrate (vertebrata_odb10) and sauropsid (sauropsida_odb10) databases. BUSCO statistics C, complete; S, single-copy; D, duplicated; F, fragmented; M, missing.

**Table S5** Summary statistics of the functionally annotated protein-coding genes

| **Database** | **#** | **%** |
| --- | --- | --- |
| CDD | 9,210 | 38 |
| Coils | 5,005 | 21 |
| GO | 20,310 | 85 |
| Gene3D | 16,390 | 35 |
| Hamap | 403 | 1 |
| InterPro | 20,310 | 44 |
| MetaCyc | 14,180 | 30 |
| MobiDBLite | 10,961 | 46 |
| PANTHER | 20,383 | 85 |
| PIRSF | 1,449 | 6 |
| PRINTS | 5,458 | 23 |
| Pfam | 18,843 | 79 |
| Phobius | 8,311 | 35 |
| ProSitePatterns | 6,252 | 26 |
| ProSiteProfiles | 11,342 | 47 |
| Reactome | 18,266 | 76 |
| SFLD | 108 | 0 |
| SMART | 10,197 | 43 |
| SUPERFAMILY | 15,477 | 65 |
| SignalPEUK | 2,826 | 12 |
| SignalPGRAMNEGATIVE | 822 | 3 |
| SignalPGRAMPOSITIVE | 1,141 | 5 |
| TIGRFAM | 1,153 | 5 |
| **Genes that have at least one hit from the databases** | 22,554 | 94.1 |
